# Plasmids persist in a microbial community by providing fitness benefit to multiple phylotypes

**DOI:** 10.1101/747964

**Authors:** Liguan Li, Arnaud Dechesne, Jonas Stenløkke Madsen, Joseph Nesme, Søren J. Sørensen, Barth F. Smets

**Author notes:** These authors contributed equally to this work (Liguan Li, Arnaud Dechesne). Corresponding author: Barth F. Smets, Address: Department of Environmental Engineering, Technical University of Denmark, Bygningstorvet, Bygning 115, 2800 Kgs. Lyngby.

## Abstract

The current epidemic of antibiotic resistance has been facilitated by the wide and rapid horizontal dissemination of antibiotic resistance genes (ARGs) in microbial communities. Indeed, ARGs are often located on plasmids, which can efficiently shuttle genes across diverse taxa. While the existence conditions of plasmids have been extensively studied in a few model bacterial populations, their fate in complex bacterial communities is poorly understood. Here, we coupled plasmid transfer assays with serial growth experiments to investigate the persistence of the broad-host-range IncP-1 plasmid pKJK5 in microbial communities derived from a sewage treatment plant. The cultivation conditions combined different nutrient and oxygen levels, and were non-selective and non-conducive for liquid-phase conjugal transfer. Following initial transfer, the plasmid persisted in almost all conditions during a 10-day serial growth experiment (equivalent to 60 generations), with a transient transconjugant incidence up to 30%. By combining cell enumeration and sorting with amplicon sequencing, we mapped plasmid fitness effects across taxa of the microbial community. Unexpected plasmid fitness benefits were observed in multiple phylotypes of *Aeromonas*, *Pseudomonas* and *Enterobacteriaceae*, which resulted in community-level plasmid persistence. We demonstrate, for the first time, that plasmid fitness effects across community members can be estimated in a high-throughput way without prior isolation. By gaining a fitness benefit when carrying plasmids, members within complex microbial communities might have a hitherto unrecognized potential to maintain plasmids for long-term community-wide access.

## Introduction

Plasmids - extrachromosomal replicons - can support rapid bacterial adaptation by moving genes between phylogenetically diverse bacteria, a process known as horizontal gene transfer [1]. This is particularly important in the case of antibiotic resistance, where acquisition of plasmid-borne antibiotic resistance genes (ARGs) can instantly render a strain impervious to antibiotic treatment [2]. The current global antibiotic resistance crisis has been largely attributed to plasmid mediated ARG dissemination [3].

Strategies to combat the antibiotic resistance epidemic therefore require an understanding of the mechanisms that underlie plasmid fate in microbial communities. Upon entering a microbial community, a plasmid will only persist if its original or secondary hosts (transconjugants) survive. If it persists, it will constitute a long-term reservoir of auxiliary genes for that community. In fact, persistence of IncP-1 plasmids or their derivatives has recently been observed in *ex situ* communities like anaerobic sludge microcosms [4] and artificial multi-species system [5] over periods longer than 10 days, as well as in murine gut microbiota experiments [6]. However, by simply interpreting plasmid persistence as the frequency of plasmid occurrence, these observational studies did not explicitly examine the mechanisms that underly plasmid persistence. Therefore, we are currently far from able to predict plasmid fate in a microbial community.

In contrast to our limited understanding of plasmid persistence at the community level, the fate of plasmids at the population level has received much more attention. There, theoretical and empirical studies have demonstrated that the conditions for plasmid persistence are governed by the interplay of segregational loss, conjugal transfer, and the effect of the plasmid on host fitness [7–9]. The rates of these processes depend on the plasmid, the host, and the environmental conditions, likely in a complex manner. First, although partitioning mechanisms exist to ensure ordered plasmid segregation prior to cell division [10], plasmid loss due to vegetative growth can typically be detected [7]. Second, plasmid carriage often imposes a reduction in fitness to the host [11]. Therefore, in the absence of mechanisms countering their segregation loss and fitness burden, plasmids would eventually be removed from bacterial populations. Plasmid persistence in a population thus requires one or both of two mechanisms: fitness cost amelioration and horizontal transfer [9].

In the majority of long-term cultivation experiments, plasmid persistence is associated with a reduction in the burden of plasmid carriage through adaptation [9]. This weakens the strength of purifying selection against plasmid carriage, and therefore reduces the rate at which plasmids are removed from the population. For example, by experimentally evolving *Pseudomonas fluorescens* carrying a high-fitness-cost plasmid, Harrison et al. observed amelioration of the cost-of-carriage across a wide gradient of environmental conditions even without positive selection for plasmid carriage [12]. Recent studies on the genetic basis underlying such adaptations among different plasmid-host pairs, suggest that poor plasmid persistence can be caused by a high cost involving helicase-plasmid interactions, which can be rapidly ameliorated [13]. Another explanation for plasmid persistence within a population is conjugal transfer. In contrast to conjugation-deficient pKJK5 derivatives, the ability of the original plasmid to conjugate played an essential role in its stable maintenance in populations of *Escherichia coli* and *Kluyvera sp*. [14,15]. More recently, Lopatkin et al. [16] experimentally demonstrated that transfer rate of common conjugative plasmids, was indeed necessary and sufficient to maintain plasmids in an *E. coli* population.

Detailed investigations of plasmid fate have been limited to pure or defined cultures of isogenic or closely related strains of a few model species. Such research on species like *P. putida* and *E. coli* have confirmed the notion that plasmid persistence is strain dependent [17–19]. Strains of *P. putida* can differ largely in their ability to maintain the IncP-1 plasmid pB10 [20,21]. In *Klebsiella pneumonia*, carrying plasmid pKP33 did not induce any burden for one strain, but the fitness of another strain was significantly reduced [19]. Yet, the cause of this inter-strain variability are currently unknown and our ability to identify them is likely limited by the fact that testing single strains is labour intensive.

Since most bacteria exist within complex communities of hundreds to thousands of species, plasmid persistence is ideally examined within this context. Population-level experiments ignore the complex interactions within communities, which prevents inferring plasmid fate in communities. For example, interspecies transfer enabled persistence of plasmid pQBR57 which would have been impossible within a single population [22]. Besides, multi-level ecological dynamics play a role in a community, since fitness differences do not only exist between plasmid-free and plasmid-bearing cells within population of singular strains, but also between strains. Therefore, a plasmid can be lost because of low intrinsic fitness of its hosts or even merely because of ecological drift.

Previous work on community permissiveness (i.e., the ability to take up a plasmid from an exogenous donor strain) has revealed that IncP-1 plasmids can readily be transferred to diverse members of complex environmental communities [23–25]. However, it is not clear to what extent the communities can maintain the plasmid and in which way the individual members can contribute to community-level plasmid persistence.

By coupling permissiveness assays using a *gfp*-tagged plasmid with serial growth experiments, we monitored the persistence of the IncP-1 plasmid pKJK5 not only at the community level but also at the level of individual phylotypes to identify the relative importance of the processes acting on plasmid persistence and how they vary across phylotypes. To reduce complexity, we chose conditions where conjugal transfer is limited and thus where segregation (in)stability, plasmid fitness effects, and competition between phylotypes would drive plasmid fate. We identified that plasmid fitness benefits in multiple phylotypes played an essential role in community-level plasmid persistence.

## Materials and Methods

### Donor strain and recipient microbial community

*E. coli* MG1655 carrying pKJK5, an IncP-1 conjugative plasmid that confers resistance to tetracycline and trimethoprim, was used as donor (further denoted *E. coli* (pKJK5)). The plasmid is marked with *gfpmut3b* under the control of a LacI^q^ repressible promoter P_*A1O4O3*_ [23]. The donor strain was chromosomally tagged with a constitutively expressed *lacI*^q^ and *mCherry* using the mini-Tn7 tagging system [23]. Therefore, plasmid-encoded *gfp* expression is repressed in the donor strain, but de-repressed following successful transfer to a recipient strain. As a result, recipients that acquire the plasmid can be quantified and retrieved as green fluorescent cells by fluorescence-activated cell sorting (FACS). The donor strain was cultured overnight in LB medium supplemented with tetracycline, harvested by centrifugation and washed in 0.9% NaCl solution. Cell density was adjusted to 3.0 × 10^7^ cells per ml by direct measurement in a Thomas counting chamber, and used in filter mating assays as described previously [26]. Activated sludge (microbial biomass) grab samples were collected from a municipal wastewater treatment plant (WWTP) (Mølleåværket, Lyngby-Taarbæk, Denmark) in March 2018, then transported refrigerated to the laboratory and processed immediately. Bacteria were extracted from 50 ml activated sludge by washing (centrifuging at approx. 5000 *g* for 10 min, carefully removing supernatant and resuspending the pellet in 0.9% NaCl) and sonication (500 J for 4 min at 50% duty cycle). After 10 min quiescent settling, 5 ml supernatant was used for counting and adjusting to 3.0 × 10^7^ cells per ml before filter mating assays.

### Filter mating assay and serial growth experiment

As the activated sludge microbial community is a mixture of enteric and non-enteric bacteria with distinct growth condition preferences, we used two media in both filter mating and serial growth experiments to selectively enrich the two community fractions (Fig. S1). MacConkey (MC) medium and synthetic wastewater (SW) [27] medium were used to favour enteric and non-enteric bacteria, respectively. In addition, we set low (100RPM) and high (500RPM) stirring rates to vary the degree of oxygen transfer during the serial growth experiment. The high stirring rate was sufficient to ensure fully oxic conditions while the low stirring rate would result in oxygen-limited to anoxic growth in the substrate rich medium (hereafter 500RPM and 100RPM are referred to as oxic and anoxic conditions). Initial plasmid transfer in the activated sludge community was facilitated via filter mating assays. Donor and recipient suspensions of 3.0 × 10^7^ cells per ml were mixed at 1:1 ratio (for matings on SW medium) or 1:5 ratio (for matings on MC medium, to compensate for the selectivity of MC media for enterobacteria), and immediately filtered onto a 0.2 μm track-etched membrane (Whatman Cyclopore^™^, UK). Filters were placed on agar-solidified (15 g/l) SW or MC medium. After 48 h incubation at 25 °C, cells from triplicate filters were combined in 2 ml 0.9% NaCl solution and detached by vortexing at 3000 RPM for 3 min. Cell number was adjusted to 3.0 × 10^7^ cells to initiate serial growth in 50 ml Erlenmeyer flasks containing 15 ml of appropriate media. The serial growth experiments were run in triplicate for 10 days under one of four combinations of media type and stirring rate (i.e., SW100, SW500, MC100 and MC500), without antibiotic selection. Daily, 1% of each culture was transferred to a new flask carrying fresh medium, ensuring growth of about 6 generations per day. Control experiments were run in parallel to assess the potential for plasmid transfer during the serial growth experiment. To this effect, donor and recipient cells were separately grown on filters (on either SW or MC medium) for 48 h at 25 °C. Then donor and recipient cells were mixed directly in flasks with liquid growth media; triplicate flasks were subjected to serial growth and treated exactly as the experimental groups under the four conditions.

### Cell sorting and 16S rRNA gene amplicon sequencing

Throughout the serial growth experiment, the relative abundance of donor, recipient and transconjugant cells were quantified daily by FACS (BD FACSAria III, USA) with the same gate settings as for sorting (see below). At selected time points during the serial growth experiment (day 2, 4, 6, 8 and 10), cell fractions were also subject to sorting and 16S rRNA gene amplicon sequencing. Transconjugant and recipient cells were sorted using FACS settings as described earlier [23]. In all sortings, we targeted a minimum of 10,000 cells. Samples with low transconjugant cell fractions (< 0.1%) were not sorted, as those conditions would require excessive time to ensure purity. Sorted cells were concentrated by centrifugation, and subject to cell lysis and DNA extraction using GenePurgeDirect^™^ DNA releasing agent (NimaGen, NL). The hypervariable regions V3-V4 of the 16S rRNA gene were PCR amplified using primer set 341F (5’-CCTAYGGGRBGCASCAG-3’) and 806R (5’-GGACTACHVGGGTWTCTAAT-3’) originally published by Yu et al [28]. and modified as described in Sundberg et al [29]. Tagging using loci-specific index primers for individual samples was done in a second PCR step. After purification using HighPrep PCR magnetic beads (MagBio, USA), sample-wise indexed PCR products were pooled in equimolar proportions and concentrated. Sequencing of pooled libraries was performed on a MiSeq platform using the v3 600 cycles sequencing kit in 300bp paired-end mode (Illumina, USA). Based on index sequences, samples were directly demultiplexed by Casava 1.8 (Illumina, USA) on the MiSeq instrument to produce the paired-end FASTQ files for sequencing analysis.

### Sequence data analysis

The DADA2 pipeline based on amplicon sequence variants (ASV) was used for 16S rRNA gene amplicon sequence analysis [30]. Briefly, filtration and trimming (maxN=0, maxEE=2, truncQ=2) were performed to clean raw sequences, which then went through error learning, de-replication and merging. By discriminating biological sequences from sequencing errors, ASVs were then inferred. Taxonomy classification of chimera-free ASVs was done by searching against the SILVA non-redundant database (release 132) [31]. The generated sample-wise ASV abundance table was used for further analysis. Each ASV is hereafter referred to as a phylotype. Only ASVs with a relative abundance threshold above 0.1% were considered for downstream analysis to limit noise attributable to rare taxa. To better place the phylogeny of *Aeromonas* phylotypes, *Aeromonas* reference 16S rRNA gene sequences were retrieved from NCBI RefSeq database, and the V3-V4 region sequence was extracted using the same primer set (341F and 806R). The phangorn R-package [32] was used to construct a phylogenetic tree, in which a neighbour-joining tree was used to fit maximum-likelihood model. Results were visualized using the R-package ggplot2 [33]. All sequences obtained in this study were deposited in NCBI under accession number PRJNA515836.

### Identification of the fate of plasmid at the phylotype level

Flow cytometry allows to quantify the density of three pools of community members and sort them: the transconjugants (cells with green fluorescence only), the recipients (cells with green fluorescence or without fluorescence), and the initial donors (cells with red fluorescence only). The absolute abundance of each phylotype in the community was estimated by multiplying the cell densities of each pool by the relative abundance of the phylotype in the transconjugant or recipient pool based on the 16S rRNA gene amplicon sequencing. Phylotypes detected at least one time point in the transconjugant pool during any serial growth experiment were defined as permissive; phylotypes never detected in the transconjugant pools were defined as non-permissive.

The overall fitness of all permissive phylotypes relative to all recipient phylotypes was calculated by examining the temporal dynamics of the ratio of their absolute abundance 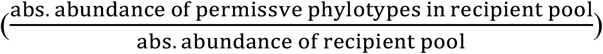 A positive or negative slope of this time-series ratio indicates an overall growth advantage or disadvantage of permissive phylotypes versus recipient phylotypes during the experiment, respectively.

For each permissive phylotype, the temporal dynamics of the fraction of plasmid-bearing cells (transconjugants) is impacted by two processes (in the absence of its *de novo* formation by horizontal gene transfer): first, the positive or negative effect of the plasmid on the phylotype’s fitness relative to plasmid-free cells of the same phylotype; second, the segregation stability or loss of the plasmid. These two processes cannot be distinguished in our experiments. To quantify the two processes, the temporal dynamics of abundance of each transconjugant phylotype relative to the same phylotype in recipient pool 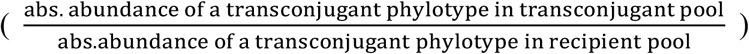 was examined. The slope (hereafter referred to as S_α+δ_) of this ratio is the combined result of plasmid segregational loss and plasmid fitness effects. In order to minimize noise and identify true positive/negative S_α+δ_, we only consider slopes biologically significant when they exceed 0.01 in absolute value.

## Results

### Plasmids persisted in microbial communities for more than 60 generations despite of negligible horizontal transfer

We created a range of environmental conditions to study plasmid persistence in complex microbial communities. The composition of the recipient pools derived from activated sludge and propagated under different conditions, became rapidly and significantly different (SW versus MC media, oxic versus anoxic) (PERMANOVA test on media and RPM effect, P-value < 0.001) (Fig. S2). The effect of mixing on community composition was the strongest in the MC medium, probably due to the oxygen-sensitivity of enterobacteria, which was also evident in the enrichment of lactose fermenting *Enterobacteriaceae* under anoxic condition compared to the oxic condition (see below). The MC medium colour changed to red under anoxic, but not under oxic conditions, confirming acid production associated with lactose fermentation.

The plasmid persisted across all examined environmental conditions, except in anoxic conditions in MC media (Fig. 1). During the first three days in the SW medium, the transconjugant density increased from < 1% in the initial filter mating to 4-8% on the 3^rd^ cultivation day. Subsequently, transconjugant densities decreased except for treatments SW100-1, SW500-2 and SW500-3, where a transient increase was observed (8-14%) before densities gradually decreased to low but stable levels (< 1%) on 10^th^ day. In the MC medium, transconjugant densities increased from 18% of initial filter mating up to 30% on the 1^st^ day. The degree of oxygen transfer had a consistent impact on transconjugant density in the MC medium, but not in the SW medium. The transconjugant densities decreased monotonically to undetectable in the anoxic (MC100) condition, while they increased transiently to 25% under oxic conditions (MC500) before gradually decreasing to low levels (< 1%).

**Figure 1.**
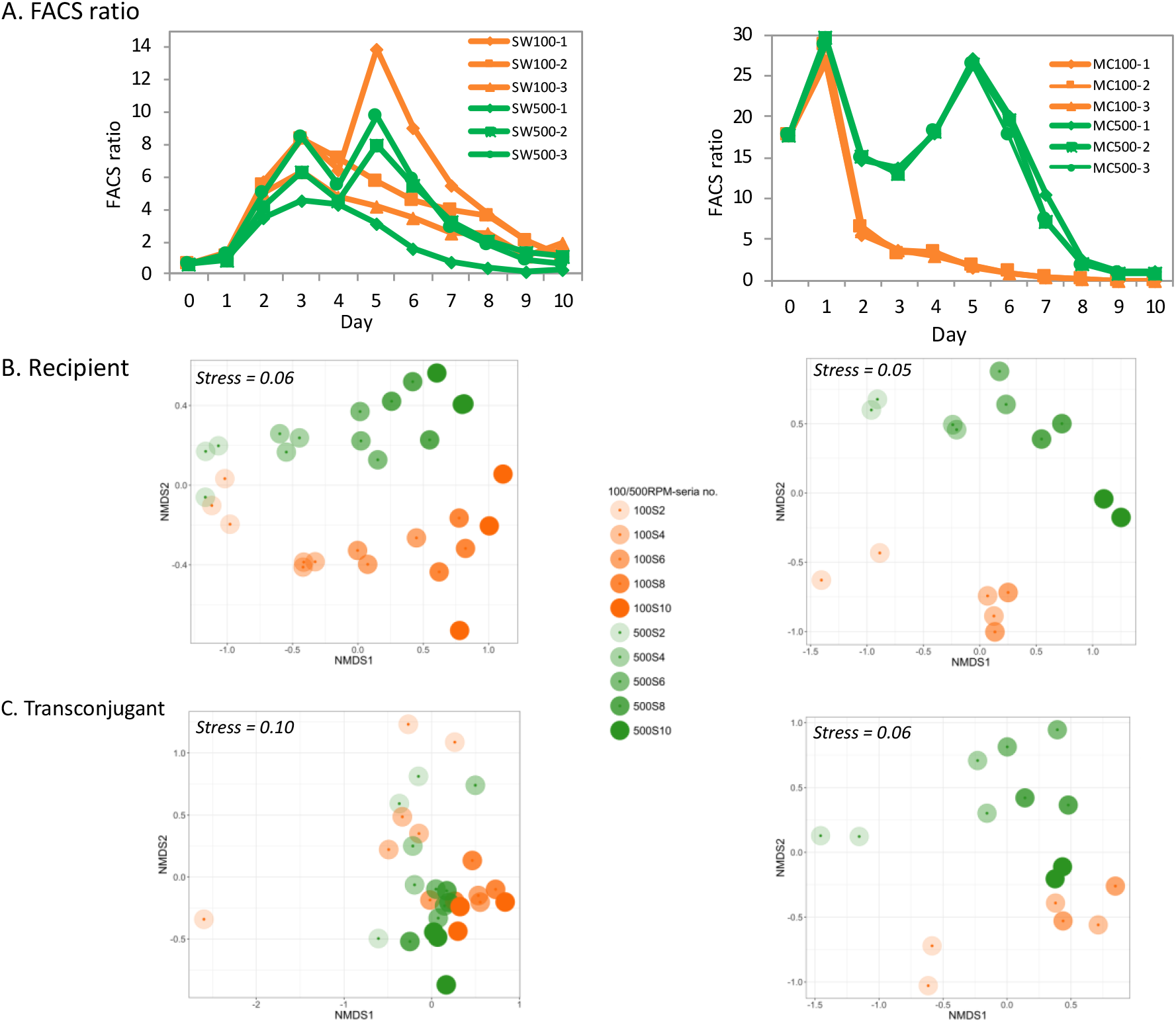
Percentage of transconjugants of a whole community and NMDS profile of microbial community composition (left: serial growth in SW; right: serial growth in MC). A. Transconjugant density detected by FACS (cells only with green fluorescence) of initial filter mating samples (0 day), daily samples through 10-day serial growth experiment (1-10 day). B&C. NMDS profile of microbial communities of recipient pools (B) and transconjugant pools (C) on 2^nd^, 4^th^, 5^th^, 6^th^, 8^th^ and 10^th^ day, stress and distance.

During the serial growth experiments, the observed transconjugant cells must derive either from new plasmid transfer events from donors to recipient cells or from persistent transconjugant cells that were formed during the initial filter mating. We included a control experiment to measure the extent of plasmid transfer during the serial growth experiment. Donor cells decreased rapidly in both control and experimental groups (e.g., from >10% on 1^st^ day to < 0.1% on 10^th^ day) (Fig. S3 and S4). Transconjugant cells were absent in almost all control groups, except for 0.1% transconjugant density in MC media on the 1^st^ day (Fig. S5). Hence, horizontal transfer of pKJK5 during the serial growth experiment was negligible.

### Persistent transconjugants spanned multiple phylotypes

While the recipient community diversity decreased through the experiment (e.g., Shannon index decreased from 4.0 to 2.0 in SW; from 3.0 to 1.7 in MC), substantial diversity remained by the end of the experiment (12 to 16 genera of 8 to 10 orders in SW; 7 to 14 genera of 5 to 6 orders in MC (Fig. 2)).

**Figure 2.**
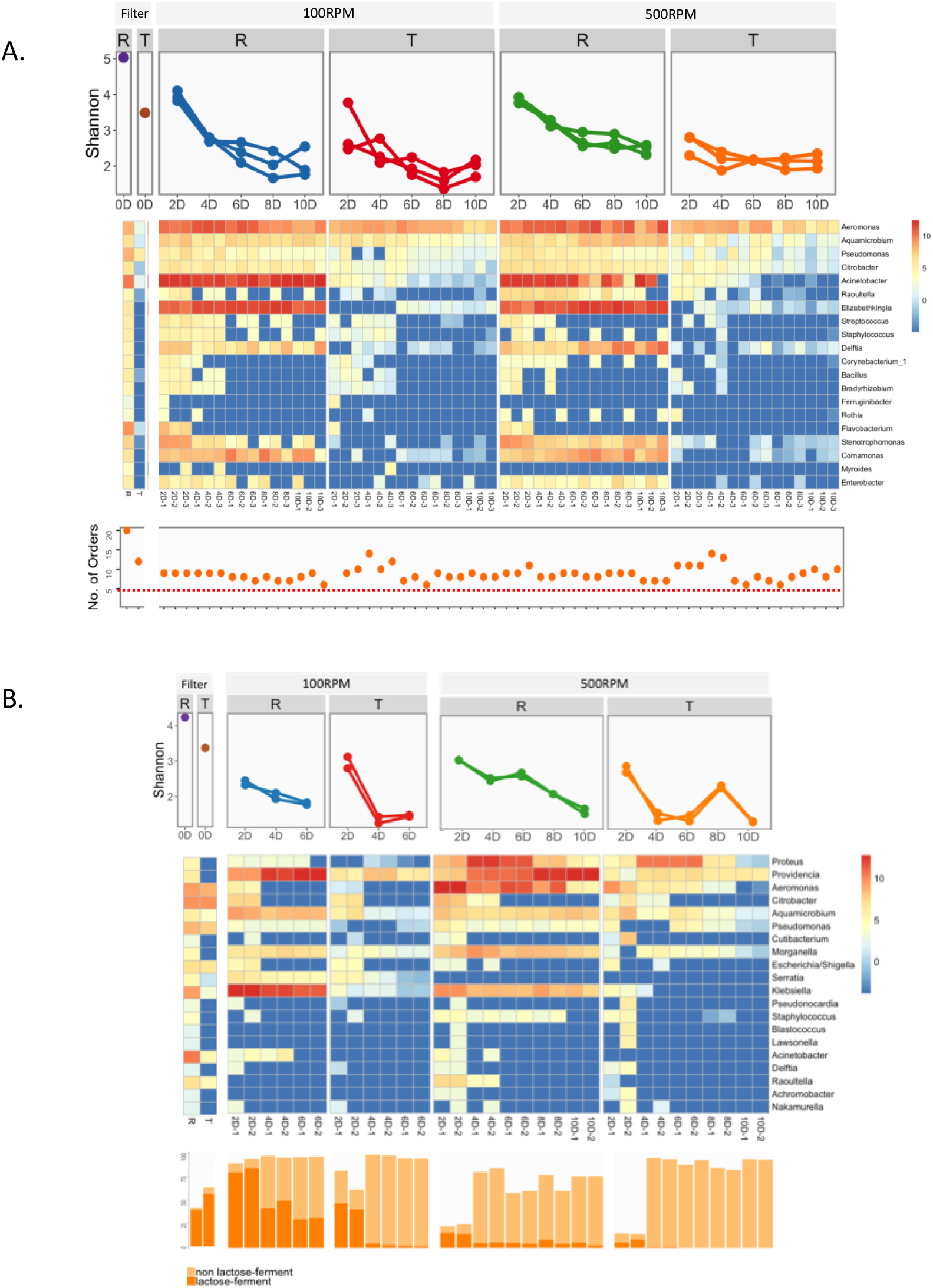
Diversity and absolute abundance of the dominant phylotypes in samples from filter mating and serial growth experiments on SW (A) and MC (B) media. Top: Shannon diversity index; middle: absolute abundance of the 20 most abundant genera; bottom: number of unique orders in SW samples (A), abundance of *Enterobacteriales* (with and without lactose fermenting ability) in MC samples (B).

During the filter mating on SW medium, pKJK5 transferred to 29 genera (4 phyla) (Fig. 2A). This diversity of transconjugants eroded gradually during serial growth in SW liquid medium (Shannon index decreased from 3.8 to 2.0). The overall composition of transconjugant pools under oxic and anoxic conditions became distinguishable (PERMANOVA, P-value = 0.027). The temporal trend of transconjugant density differed among series possibly caused by distinct dynamic profiles among individual phylotypes. We indeed identified multiple phylotypes with absolute abundance highly correlated with the transconjugant density profile during serial growth experiment (r > 0.9, Pearson correlation), e.g., phylotypes in genera of *Citrobacter* and *Enterobacter* in SW100-1, *Stenotrophomonas* and *Aeromonas* in SW500-2 (Fig. S6). After 10 days, 62 phylotypes in 9 to14 genera were detected across the transconjugant pools. A core of 12 persistent phylotypes in 5 genera (*Aeromonas*, *Aquamicrobium*, *Citrobacter*, *Elizabethkingia* and *Pseudomonas*) were shared among all SW experiments (Fig. 3A). In each transconjugant pool, *Aeromonas* phylotypes became enriched (relative abundance > 50%) (Fig. S7). In total, 37 different *Aeromonas* phylotypes were detected in the recipient pools grown on SW, 12 of which persisted as transconjugants throughout at least one serial growth experiment (Fig. S8). These *Aeromonas* phylotypes spanned multiple species (Fig. S9), where the dominant ASV6 and ASV1 are closely related to the human opportunistic pathogens *A. veronii* and *A. hydrophila* [34], respectively.

**Figure 3.**
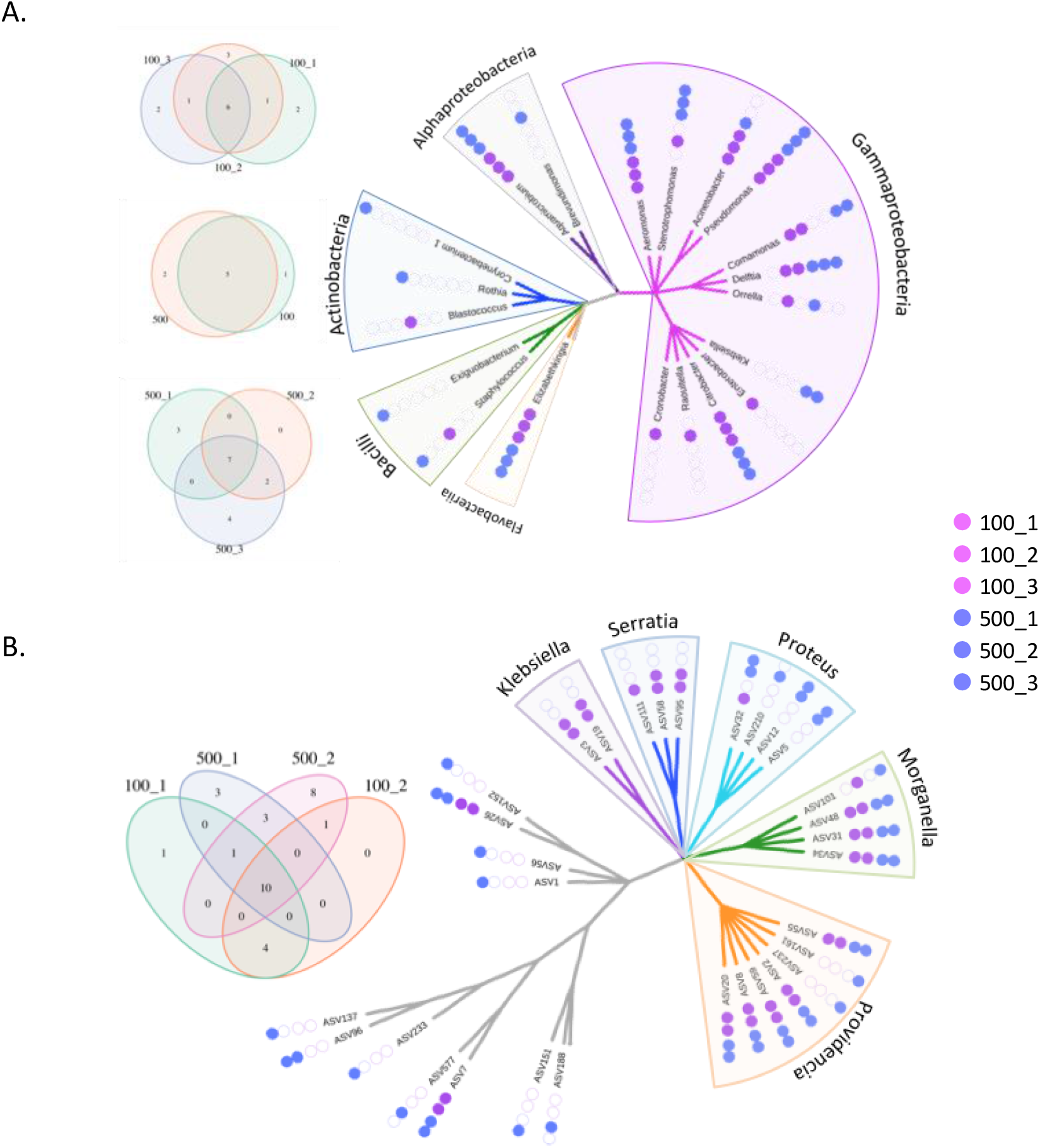
Persistent genera in SW (A) and persistent phylotypes in MC (B) at the end of the serial growth experiment. Given the higher diversity observed in SW compared to MC, genus level results are presented for SW and phylotype level results for MC. Venn diagrams show shared and unique genera/phylotypes among groups. Phylogenetic trees show all persistent genera/phylotypes. Class names are used as label for SW, while genus names in *Enterobacteriales* order are used for MC.

In MC medium, both transconjugant and recipient pools differed significantly from those that developed in SW medium during the 10-day experiment (Fig. 2B and Fig. S2). On MC medium, pKJK5 was initially transferred to 18 genera (2 phyla) in filter matings, mainly consisting of genera *Citrobacter* (48% relative abundance), *Aeromonas* (23%), and *Pseudomonas* (12%). In the serial growth experiment, transconjugant diversity continuously decreased, indicated by Shannon index decrease from 3.0 to 1.0. Oxygen conditions significantly affected the composition of transconjugant pools (PERMANOVA, P-value < 0.001). Several phylotypes had abundances that were strongly correlated (r > 0.9, Pearson correlation) with the observed dynamic profile of transconjugant density, such as *Proteus* and *Providencia* under oxic condition (Fig. S6). Transconjugants of the *Enterobacteriaceae* family were more dominant under anoxic than oxic condition (89-96% and 56-98% versus 24-84% and 13-95% in recipient and transconjugant pools, respectively). Especially, in the transconjugant pool, genera of lactose-fermenting enterobacterial species (i.e., *Escherichia*, *Enterobacter* and *Klebsiella*) were transiently abundant under anoxic conditions (up to 47%), but they were always rare under oxic conditions (< 9%). At the end of the experiment, 31 transconjugant phylotypes in 13 genera were detected, in which 8 core phylotypes in *Enterobacteriales* were shared across all treatments, including 3 *Morganella* phylotypes and 5 *Providencia* phylotypes (Fig. 3B).

### No general fitness deficit of permissive phylotypes relative to the rest of the community

We noted an average 70% decrease in the richness of permissive phylotypes during all serial growth experiments (Fig. S10). In other words, (potential) plasmid hosts disappeared from the community because of insufficient fitness or ecological drift. For example, several *Acinetobacter* and *Pseudomonas* phylotypes, abundant at the start of the serial growth experiment in SW media, progressively and simultaneously became absent from both recipient and transconjugant pools. This process would have limited plasmid persistence if permissive phylotypes were lost faster than the rest of the community. The only condition where this might have been the case was MC100 (Fig. 4), although the slope is not significantly different from 0 because only 3 data points are available. MC100 was indeed the condition where the plasmid failed to persist. In this case, strong selection against some permissive phylotypes contributed to the disappearance of transconjugants and thus to plasmid loss. Elsewhere, the permissive phylotypes appeared to decline equally or less rapidly than the rest of the community, with a similar effect of richness loss on recipient and transconjugant pools within a community thus a neutral or positive effect on plasmid persistence.

**Figure 4.**
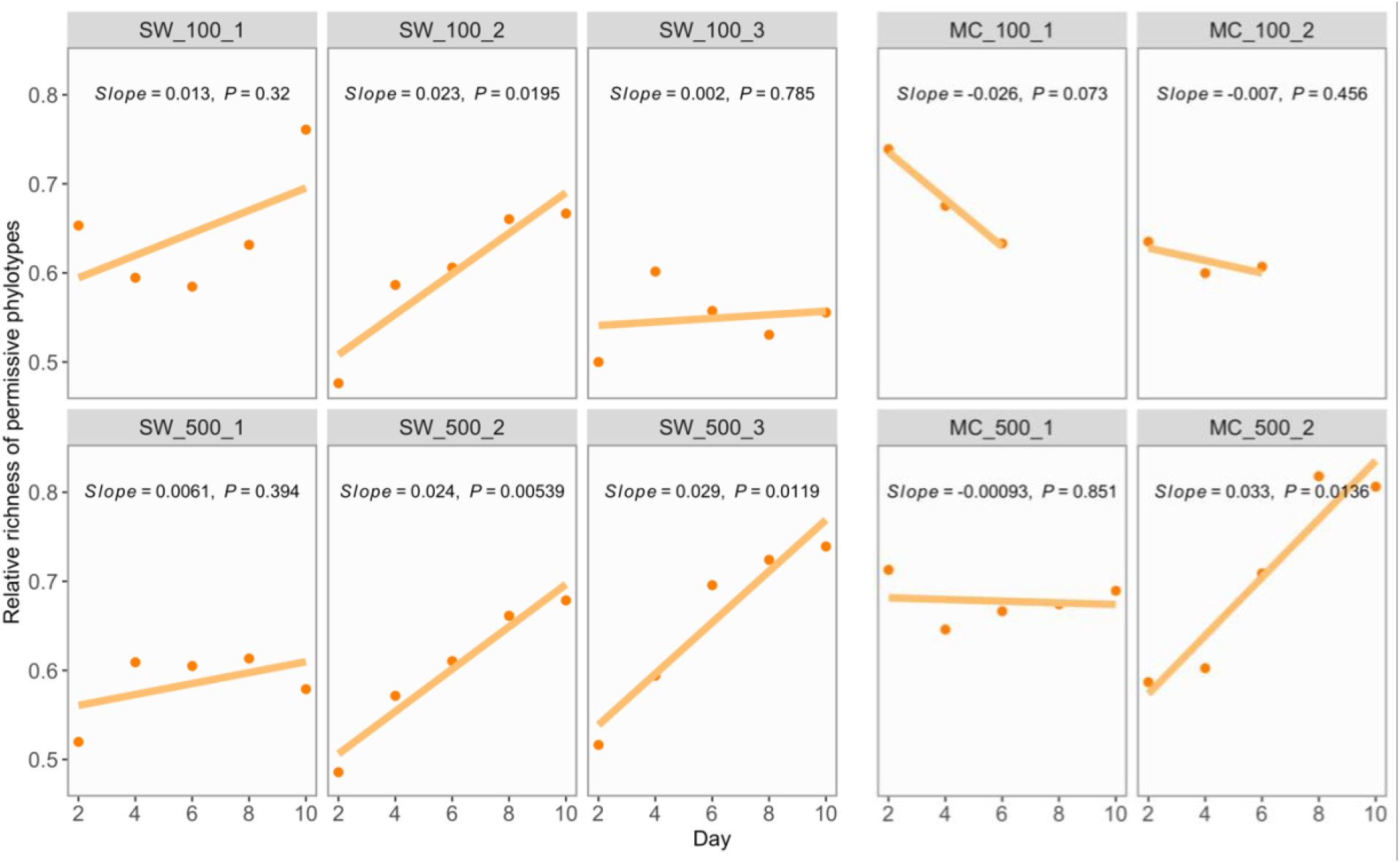
Richness of permissive phylotypes relative to all recipient phylotypes over time series. A linear regression model was applied to estimate the temporal trend of the relative richness. Solid lines represent regression lines labelled with slope and P-value. Positive or negative slope indicated slower or faster richness loss trends in permissive phylotypes compared to recipient phylotypes.

As removal from the community is an extreme outcome of low fitness, we also compared average fitness of permissive and non-permissive phylotypes to look for an overall fitness differential. Across all experimental conditions, the average growth rate of permissive phylotypes was similar or higher compared to the rest of the community (no significantly negative slopes in Fig. S11). Thus, we did not detect selection against the permissive fraction of the community. In fact, permissive phylotypes in *Aeromonas*, *Elizabethkingia* and *Acinetobacter* had a clear growth advantage, as their relative abundance increased from 30% to 80% through the serial growth experiment, which provided potential for plasmid maintenance.

### Fitness benefit associated with plasmid carriage improved persistence in multiple phylotypes

The presence of the same phylotype in the plasmid-free and plasmid-carrying fractions of the community gave us the opportunity to assess their relative dynamics and explore the cumulated effect (denoted as S_α+δ_) of plasmid segregation stability/loss and plasmid-induced fitness benefit/burden at the phylotype level at high-throughput (Fig. 5). A positive value of S_α+δ_ indicates a net fitness benefit associated with plasmid carriage since the contribution of plasmid segregation effect can only be neutral or negative. In contrast, a zero to negative value of S_α+δ_ makes it difficult to assess the strength and direction of the plasmid fitness effect because plasmid fitness burden/benefit and plasmid segregational stability/loss are indistinguishable in this study (e.g., S_α+δ_ = 0 can be caused by either (segregational loss + fitness benefit) or (segregational stability + no fitness effect)).

**Figure 5.**
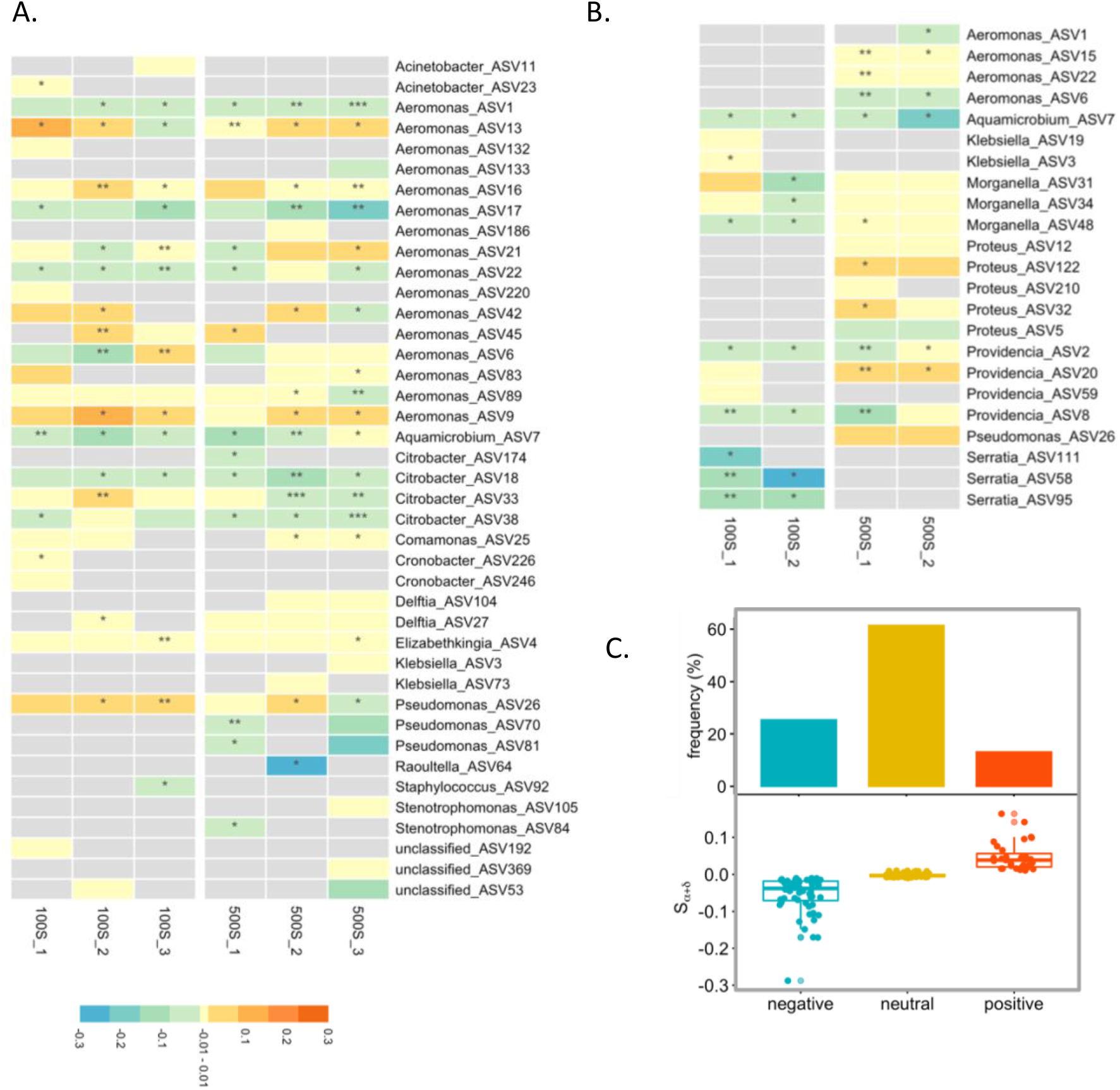
S_α+δ_ value profile in SW media (A), MC media (B) and all experimental groups (C). S_α+δ_ is an estimate of the combined plasmid fitness and segregation effects. −0.01≤ S_α+δ_ ≤ 0.01 or S_α+δ_ < −0.01: neutral or negative plasmid effect; S_α+δ_ > 0.01: positive plasmid effect. P-values of slope: * < 0.5; ** < 0.1; *** < 0.01. Occurrence frequency of three S_α+δ_ group with negative, neutral and positive values in (A) and (B) are summarized in (C).

Across different experimental conditions, we identified diverse S_α+δ_ profiles with both positive and negative values among transconjugant phylotypes (hereafter a phylotype under one specific experimental condition is indicated as ASV_expt_, e.g., ASV1_SW100-1_) (Fig. 5A&B). In SW media more than half of the ASV_expt_s were without any negative plasmid effect (79 out of 132 ASV_expt_s with S_α+δ_ ≥ 0, of which 25 with S_α+δ_ > 0). Especially, within the genus *Aeromonas*, there was no negative plasmid effect for 40 of 67 ASV_expt_s (S_α+δ_ ≥ 0), with obvious fitness benefit for 20 ASV_expt_s (S_α+δ_ > 0). In particular, three *Aeromonas* phylotypes (ASV9, ASV13 and ASV42) showed fitness benefits in the majority of the serial growth experiments. In other genera like *Pseudomonas, Delftia* and *Elizabethkingia*, there were also a few phylotypes with neutral or even positive plasmid effects. In SW media, the plasmid had a similar effect on phylotypes irrespective of oxygen transfer conditions. However, in MC media, a neutral or positive plasmid effect was more often detected under oxic than anoxic conditions, e.g., 72% (22 out of 32) and 28% (6 out of 21) of ASV_expt_s with S_α+δ_ ≥ 0 under oxic and anoxic conditions respectively. We identified 28 out of 53 ASV_expt_s with neutral (20 with S_α+δ_ = 0) or positive (8 with S_α+δ_ > 0) plasmid effects in MC media. Overall, across experimental conditions in both SW and MC media, more than half (107 out of 185) of the examined ASV_expt_s maintained the plasmid without any negative plasmid effect, even several (33 out of 185) with fitness benefit (Fig. 5C).

## Discussion

We examined, for the first time, a complex environmental community to quantitatively monitor persistence of an IncP-1 plasmid pKJK5 among diverse taxa in serial growth experiments. Without any antibiotic pressure to select for transconjugants, we focused on the potential of the community to maintain the plasmid under different nutrient and oxygen conditions. We could discount the contribution of new transfer events to the plasmid fate, as supported by the negligible transfer of pKJK5 in the control experiments and consistent with the previously reported low transfer of pKJK5 in liquid conditions [15]. Indeed, Plasmids (IncP/N/W) with short and rigid pili only transfer efficiently on solid surfaces, unlike those with long and flexible pili (IncF/H/T/J), capable of transferring equally well in liquid and on solid surfaces [35,36].

We experimentally demonstrate community-level plasmid persistence across different environmental conditions in the absence of intentional selection for plasmid carriage and in the absence of significant *de novo* transfer. In SW and MC media, time-series profiles of transconjugant density and composition became significantly different. The role of the environment in shaping transconjugant composition became obvious, especially under oxic and anoxic conditions in MC media. In fact, earlier observations have consistently identified the importance of the environment in determining plasmid fate, by either setting up microcosm under different oxygen conditions (e.g., aerobic and anaerobic sludge reactors) [4] or manipulating other environmental factors (e.g., spatial structure or presence/absence of antibiotics) [5].

By examining plasmid effects temporally across taxa in complex microbial communities, we dissected processes contributing to plasmid persistence within a community. We observed that a plasmid fitness benefit in multiple phylotypes contributed strongly to plasmid persistence at the community level. We expected plasmid pKJK5 to be a neutral or even detrimental extra-chromosomal element for the fitness of its host, as the plasmid does not encode any apparent beneficial phenotypes. Although plasmid fitness cost amelioration are frequently observed in population-level studies over evolutionary time scale (> 500 generations), it can also appear as early as within 50 generations, such as the drastically improved stability of IncP-1 plasmid pBP136 in *E. coli* under non-selective conditions [37]. Out of those experiments on plasmid-host compensatory coevolution, genomic changes frequently emerged within 100 generations [19,38]. Therefore, early genomic adaptations might have occurred improving plasmid fitness and persistence within the time-scale of the experiment. It is also possible that the microbial community we used as potential recipients might already have been subjected to compensatory adaptation for hosting IncP-1 plasmids. Genomic and experimental studies have demonstrated a high prevalence of IncP-1 plasmids spanning all subgroups (IncP-1 α, β, γ, δ and ε) in WWTP systems [39–41]. If this were the case, community members already experiencing genetic interaction with IncP-1 plasmids might favour carrying cognate plasmids like pKJK5. If this could explain why pKJK5 might have caused minimal burden to some phylotypes, it does not explain why it would provide a fitness benefit. Further experiments with specific knockout mutant of the plasmid genes will likely be necessary to provide a molecular explanation.

The wide range of S_α+δ_ values across phylotypes and conditions confirmed that the plasmid effect is a phylotype and context dependent feature. In fact, population-level studies have repeatedly revealed that plasmid effects are indeed determined by strain-specific factors [42,43]. We identify here, for the first time, individual specific plasmid effect within a complex microbial community. Although the underlying mechanisms are not yet clear, the heterogenous plasmid effect observed in this study might be the overall result of phylotype-level specific compensatory adaptations, protective mechanisms, and/or regulatory networks. For example, as revealed in plasmid-host interaction in clinical isolates of *K. pneumoniae* and *E. coli* [19], differential adaptive evolutions among strains and species modifies the plasmid effect on hosts in various ways.

Across the experimental conditions, plasmid persistence was especially observed in several phylotypes in the *Aeromonas, Pseudomonas, Proteus* and *Providencia* genera. In particular, several phylotypes of *Aeromonas* received fitness benefits from pKJK5 carriage under different experimental conditions. These genera have often been reported as carriers of antibiotic resistance plasmids in either environmental or clinical settings, such as plasmid mediated carbapenem- and colistin-resistance in *Proteus* and *Providencia* isolates. Recent studies have demonstrated the high potential of *Aeromonas* in carrying multidrug-resistance plasmids [44–46], and in transferring plasmids in various environmental communities [23–25]. Given the ubiquitous presence of *Aeromonas* in aquatic environments [47,48], *Aeromonas* might play a significant role in facilitating transfer and maintenance of plasmid-mediated ARGs in environmental communities.

We demonstrated that complex microbial communities can maintain plasmids under non-selective conditions, and phylotype-level plasmid fitness benefits significantly facilitate community-level persistence. To further understand the role of the community in modulating plasmid behaviour, the distinct mechanisms of plasmid-host interactions in communities compared with populations remain to be elucidated.

## Acknowledgements

This work was supported by the Joint Programming Initiative-Antimicrobial Resistance (JPI-AMR; DARWIN project #7044-00004B) to BFS; and the H.C. Ørsted Postdoc programme co-funded by Marie Skłodowska-Curie Actions to LL.

## References

1. Soucy SM, Huang J, Gogarten JP. Horizontal gene transfer: building the web of life. Nat Rev Genet. 2015;16:472–82.

2. Holmes AH, Moore LSP, Sundsfjord A, Steinbakk M, Regmi S, Karkey A, et al. Understanding the mechanisms and drivers of antimicrobial resistance. Lancet. 2016;387:176–87.

3. Crofts TS, Gasparrini AJ, Dantas G. Next-generation approaches to understand and combat the antibiotic resistome. Nat Rev Microbiol. 2017;15:422–34.

4. Merlin C, Bonot S, Courtois S, Block JC. Persistence and dissemination of the multiple-antibiotic-resistance plasmid pB10 in the microbial communities of wastewater sludge microcosms. Water Res. 2011;45:2897–905.

5. Cairns J, Ruokolainen L, Hultman J, Tamminen M, Virta M, Hiltunen T. Ecology determines how low antibiotic concentration impacts community composition and horizontal transfer of resistance genes. Commun Biol. 2018;1–8.

6. Ronda C, Chen SP, Cabral V, Yaung SJ, Wang HH. Metagenomic engineering of the mammalian gut microbiome in situ. Nat Methods. 2019;16:167–70.

7. Bergstrom CT, Lipsitch M, Levin BR. Natural selection, infectious transfer and the existence conditions for bacterial plasmids. Genetics. 2000;155:1505–19.

8. Levin BR, Stewart FM. The population biology of bacterial plasmids: a priori conditions for the existence of mobilizable nonconjugative factors. Genetics. 1980;94:425–43.

9. Harrison E, Brockhurst MA. Plasmid-mediated horizontal gene transfer is a coevolutionary process. Trends Microbiol. 2012;20:262–7.

10. Pinto UM, Pappas KM, Winans SC. The ABCs of plasmid replication and segregation. Nat Rev Microbiol. 2012;10:755–65.

11. San Millan A, MacLean RC. Fitness costs of plasmids: a limit to plasmid transmission. Microbiol Spectr. 2017;5:1–12.

12. Harrison E, Guymer D, Spiers AJ, Paterson S, Brockhurst MA. Parallel compensatory evolution stabilizes plasmids across the parasitism-mutualism continuum. Curr Biol. 2015;25:2034–9.

13. Loftie-Eaton W, Bashford K, Quinn H, Dong K, Millstein J, Hunter S, et al. Compensatory mutations improve general permissiveness to antibiotic resistance plasmids. Nat Ecol Evol. 2017;1:1354–63.

14. Lundquist PD, Levin BR. Transitory derepression and the maintenance of conjugative plasmids. Genetics. 1986;113:483–97.

15. Bahl MI, Hansen LH, Sørensen SJ. Impact of conjugal transfer on the stability of IncP-1 plasmid pKJK5 in bacterial populations. FEMS Microbiol Lett. 2007;266:250–6.

16. Lopatkin AJ, Meredith HR, Srimani JK, Pfeiffer C, Durrett R, You L. Persistence and reversal of plasmid-mediated antibiotic resistance. Nat Commun. 2017;8:1689.

17. Hall JPJ, Harrison E, Lilley AK, Paterson S, Spiers AJ, Brockhurst MA. Environmentally co-occurring mercury resistance plasmids are genetically and phenotypically diverse and confer variable context-dependent fitness effects. Environ Microbiol. 2015;17:5008–22.

18. Heuer H, Ebers J, Weinert N, Smalla K. Variation in permissiveness for broad-host-range plasmids among genetically indistinguishable isolates of *Dickeya* sp. from a small field plot. FEMS Microbiol Ecol. 2010;73:190–6.

19. Porse A, Schønning K, Munck C, Sommer MOA. Survival and evolution of a large multidrug resistance plasmid in new clinical bacterial hosts. Mol Biol Evol. 2016;33:2860–73.

20. Heuer H, Fox RE, Top EM. Frequent conjugative transfer accelerates adaptation of a broad-host-range plasmid to an unfavorable *Pseudomonas putida* host. FEMS Microbiol Ecol. 2007;59:738–48.

21. De Gelder L, Ponciano JM, Joyce P, Top EM. Stability of a promiscuous plasmid in different hosts: No guarantee for a long-term relationship. Microbiology. 2007;153:452–63.

22. Hall JPJ, Wood AJ, Harrison E, Brockhurst MA. Source–sink plasmid transfer dynamics maintain gene mobility in soil bacterial communities. Proc Natl Acad Sci. 2016;113:8260–5.

23. Klümper U, Riber L, Dechesne A, Sannazzarro A, Hansen LH, Sørensen SJ, et al. Broad host range plasmids can invade an unexpectedly diverse fraction of a soil bacterial community. ISME J. 2015;9:934–45.

24. Li L, Dechesne A, He Z, Madsen JS, Nesme J, Sørensen SJ, et al. Estimating the transfer range of plasmids encoding antimicrobial resistance in a wastewater treatment plant microbial community. Environ Sci Technol Lett. 2018;5:260–5.

25. Jacquiod S, Brejnrod A, Morberg SM, Abu Al-Soud W, Sørensen SJ, Riber L. Deciphering conjugative plasmid permissiveness in wastewater microbiomes. Mol Ecol. 2017;26:3556–71.

26. Klümper U, Dechesne A, Smets BF. Protocol for evaluating the permissiveness of bacterial communities toward conjugal plasmids by quantification and isolation of transconjugants. In: McGenity TJ, Timmis KN, Balbina N, editors. Hydrocarbon and Lipid Microbiology Protocols: Genetic, Genomic and System Analyses of Communities. Springer Berlin Heidelberg; 2014. p. 275–88.

27. Test No. 303: Simulation Test - Aerobic Sewage Treatment. In: OECD Guidelines for the Testing of Chemicals, Section 3. OECD Publishing; 2013. p. 50.

28. Yu Y, Lee C, Kim J, Hwang S. Group-specific primer and probe sets to detect methanogenic communities using quantitative real-time polymerase chain reaction. Biotechnol Bioeng. 2005;89:670–9.

29. Sundberg C, Al-Soud WA, Larsson M, Alm E, Yekta SS, Svensson BH, et al. 454 pyrosequencing analyses of bacterial and archaeal richness in 21 full-scale biogas digesters. FEMS Microbiol Ecol. 2013;85:612–26.

30. Callahan BJ, McMurdie PJ, Rosen MJ, Han AW, Johnson AJA, Holmes SP. DADA2: High-resolution sample inference from Illumina amplicon data. Nat Methods. 2016;13:581–3.

31. Quast C, Pruesse E, Yilmaz P, Gerken J, Schweer T, Yarza P, et al. The SILVA ribosomal RNA gene database project: Improved data processing and web-based tools. Nucleic Acids Res. 2013;41:590–6.

32. Schliep KP. phangorn: Phylogenetic analysis in R. Bioinformatics. 2011;27:592–3.

33. Wickham H. ggplot2 - elegant graphics for data analysis (2nd edition). New York, NY: Springer; 2016.

34. Janda JM, Abbott SL. The genus Aeromonas: Taxonomy, pathogenicity, and infection. Clin Microbiol Rev. 2010;23:35–73.

35. Bradley DE. Determination of pili by conjugative bacterial drug resistance plasmids of incompatibility groups B, C, H, J, K, M, V, and X. J Bacteriol. 1980;141:828–37.

36. Bradley DE. Characteristics and function of thick and thin conjugative pili determined by transfer-derepressed plasmids of incompatibility groups I1, I2, I5, B, K and Z. Microbiology. 1984;130:1489–502.

37. Sota M, Top EM. Host-specific factors determine the persistence of IncP-1 plasmids. World J Microbiol Biotechnol. 2008;24:1951–4.

38. Bottery MJ, Wood AJ, Brockhurst MA. Temporal dynamics of bacteria-plasmid coevolution under antibiotic selection. ISME J. 2018;13:559–562.

39. Zhang T, Zhang X-X, Ye L. Plasmid metagenome reveals high levels of antibiotic resistance genes and mobile genetic elements in activated sludge. PLoS One. 2011;6:e26041.

40. Schlüter A, Szczepanowski R, Pühler A, Top EM. Genomics of IncP-1 antibiotic resistance plasmids isolated from wastewater treatment plants provides evidence for a widely accessible drug resistance gene pool. FEMS Microbiol Rev. 2007;31:449–77.

41. Bahl MI, Burmølle M, Meisner A, Hansen LH, Sørensen SJ. All IncP-1 plasmid subgroups, including the novel ε subgroup, are prevalent in the influent of a Danish wastewater treatment plant. Plasmid. 2009;62:134–9.

42. Vogwill T, Maclean RC. The genetic basis of the fitness costs of antimicrobial resistance: a meta-analysis approach. Evol Appl. 2015;8:284–295.

43. Kottara A, Hall JPJ, Harrison E, Brockhurst MA. Variable plasmid fitness effects and mobile genetic element dynamics across *Pseudomonas* species. FEMS Microbiol Ecol. 2018;94:1–7.

44. Olaniran AO, Nzimande SBT, Mkize NG. Antimicrobial resistance and virulence signatures of *Listeria* and *Aeromonas* species recovered from treated wastewater effluent and receiving surface water in Durban, South Africa. BMC Microbiol. 2015;15:234.

45. Popowska M. Occurrence and variety of β-lactamase genes among *Aeromonas* spp. isolated from urban wastewater treatment plant. Front Microbiol. 2017;8:1–12.

46. Hedges R. W. PS, Brazil G. Resistance plasmids of *Aeromonads*. J Gen Microbiol. 1985;2091–5.

47. Piotrowska M, Popowska M. The prevalence of antibiotic resistance genes among Aeromonas species in aquatic environments. 2014;921–34.

48. Ye L, Zhang T. Pathogenic bacteria in sewage treatment plants as revealed by 454 pyrosequencing. Environ Sci Technol. 2011;45:7173–9.

