## Supplementary information for "Plasmids persist in a microbial community by providing fitness benefit to multiple phylotypes"

^*^Corresponding author

Figures

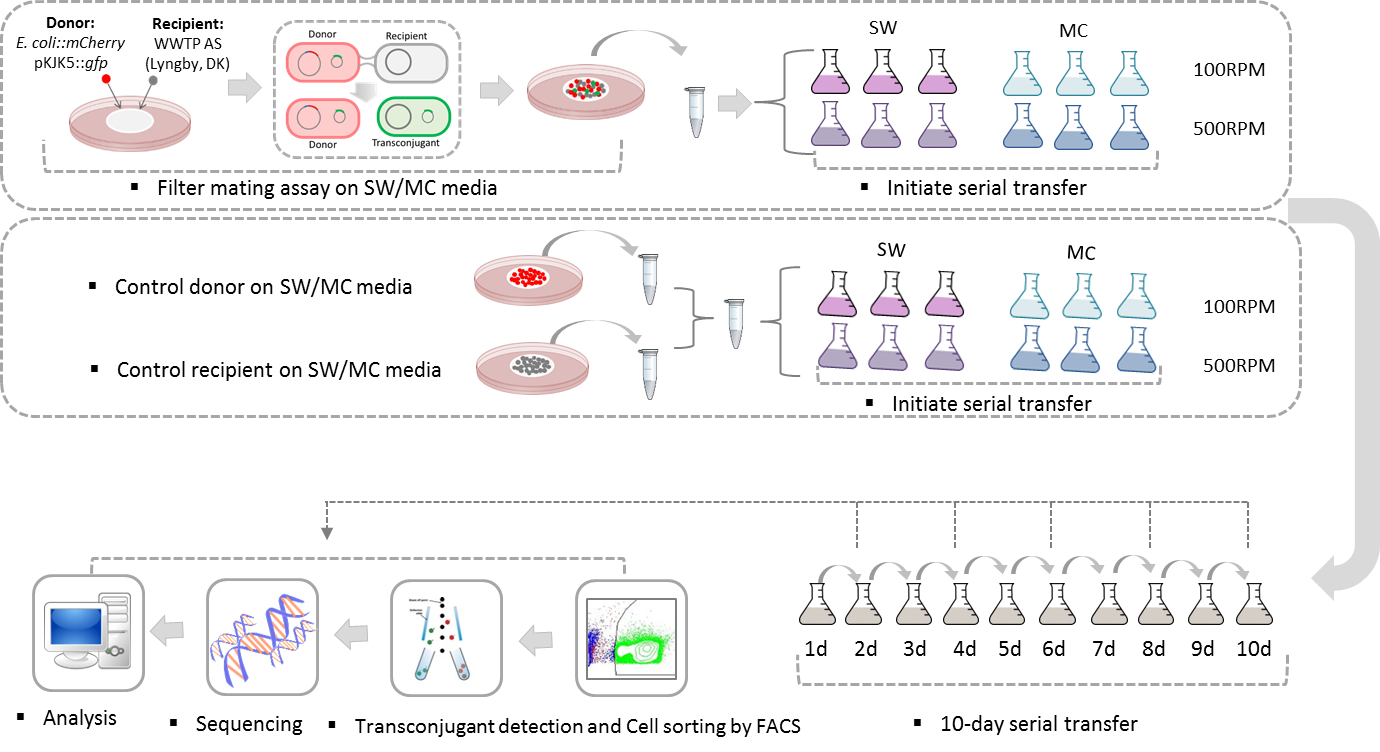

Figure S1. Experimental design of the whole serial transfer experiment (including control experiment).

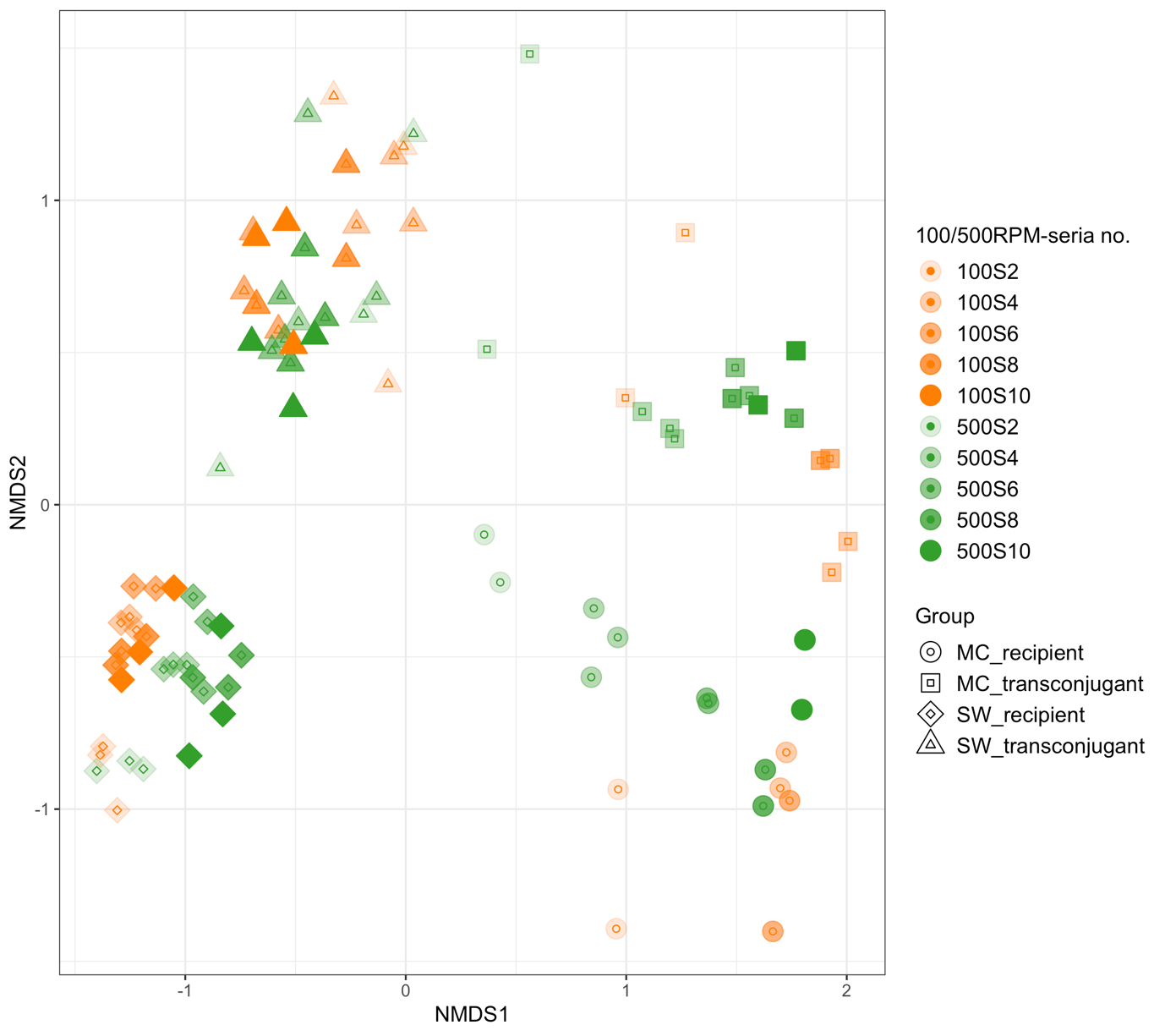

Figure S2. NMDS profile of community composition of all recipient and transconjugant pools.

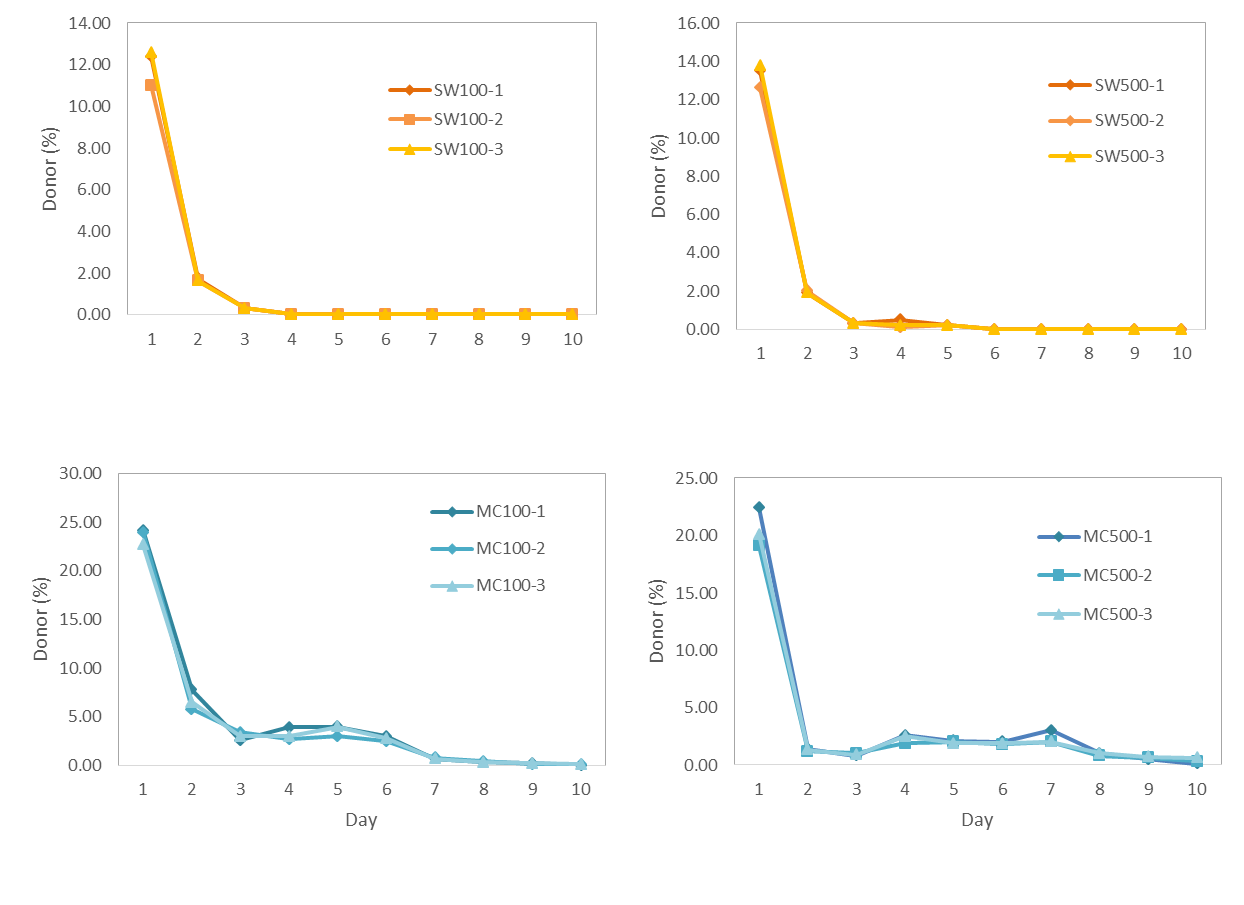

Figure S3. Relative abundance of the donor bacterium *E. coli* MG1665 (pKJK5) in control groups where donor and recipient community where co-inoculated on the first day without prior filter mating.

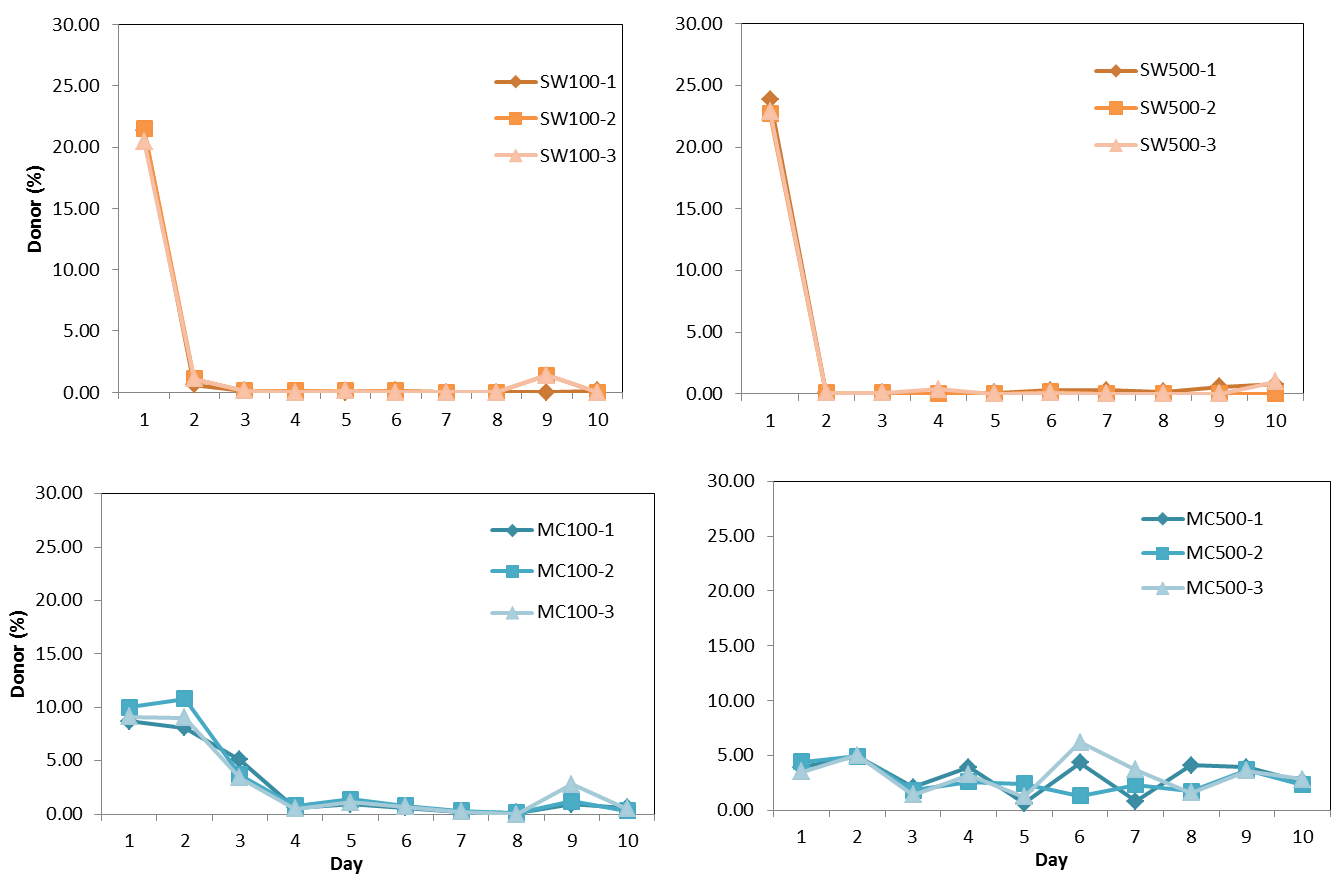

Figure S4. Relative abundance of the donor bacterium *E. coli* MG1665 (pKJK5) in experimental groups where serial growth experiments were initiated by filter mating.

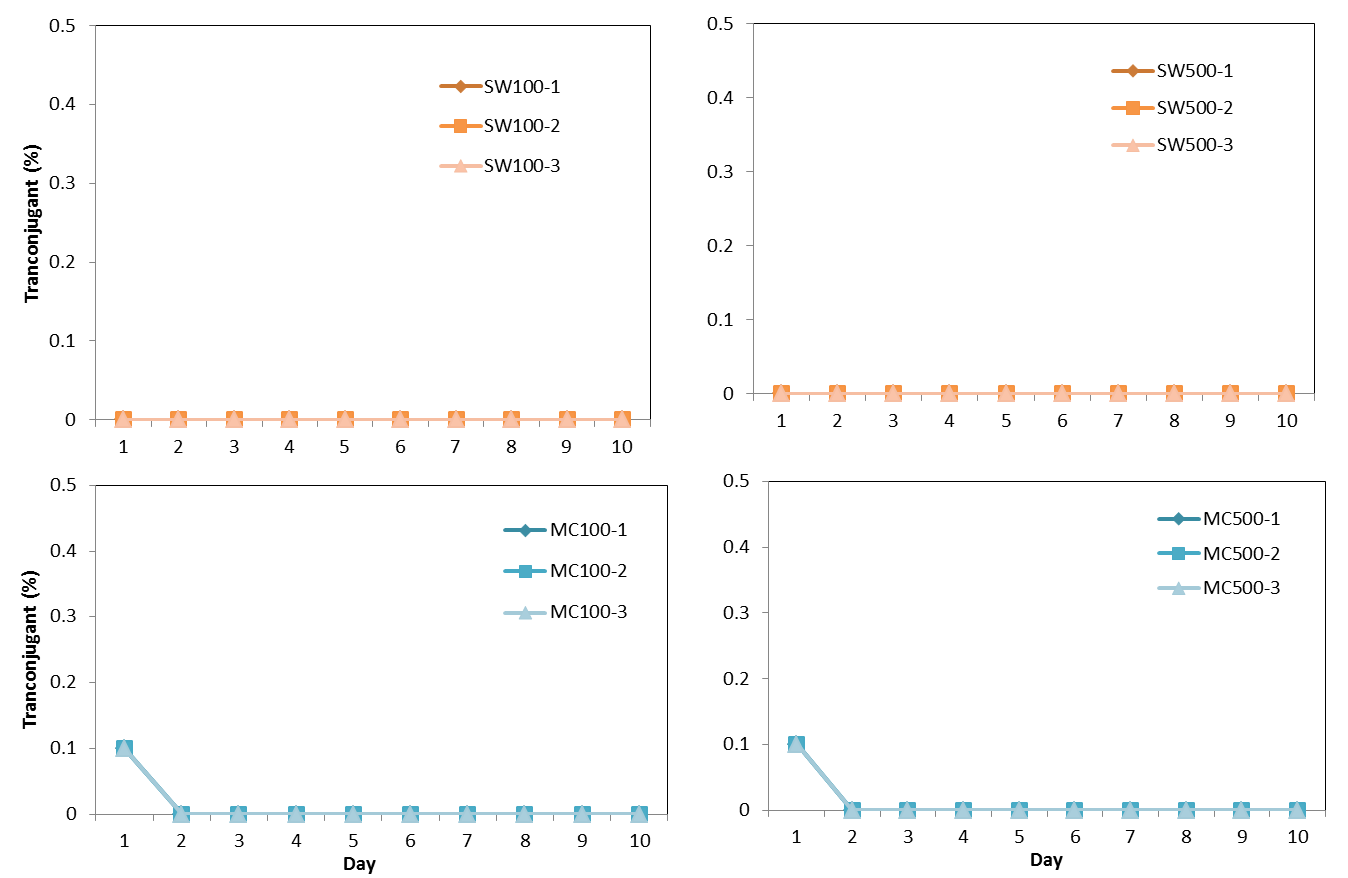

Figure S5. Relative abundance of the transconjugant in control groups where donor and recipient community where co-inoculated on the first day without prior filter mating.

A. SW medium

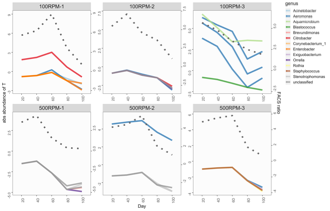

B. MC medium

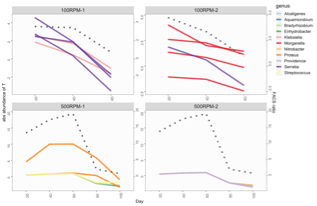

Figure S6. Temporal dynamics of the 5 phylotypes with absolute abundance most highly correlated with temporal dynamic profile of transconjugant density during serial growth experiments in SW (A) and MC media (B). The left Y-axis is absolute abundance (log2 transformed) of the phylotypes (solid line), right Y-axis is transconjugant density (dotted line).

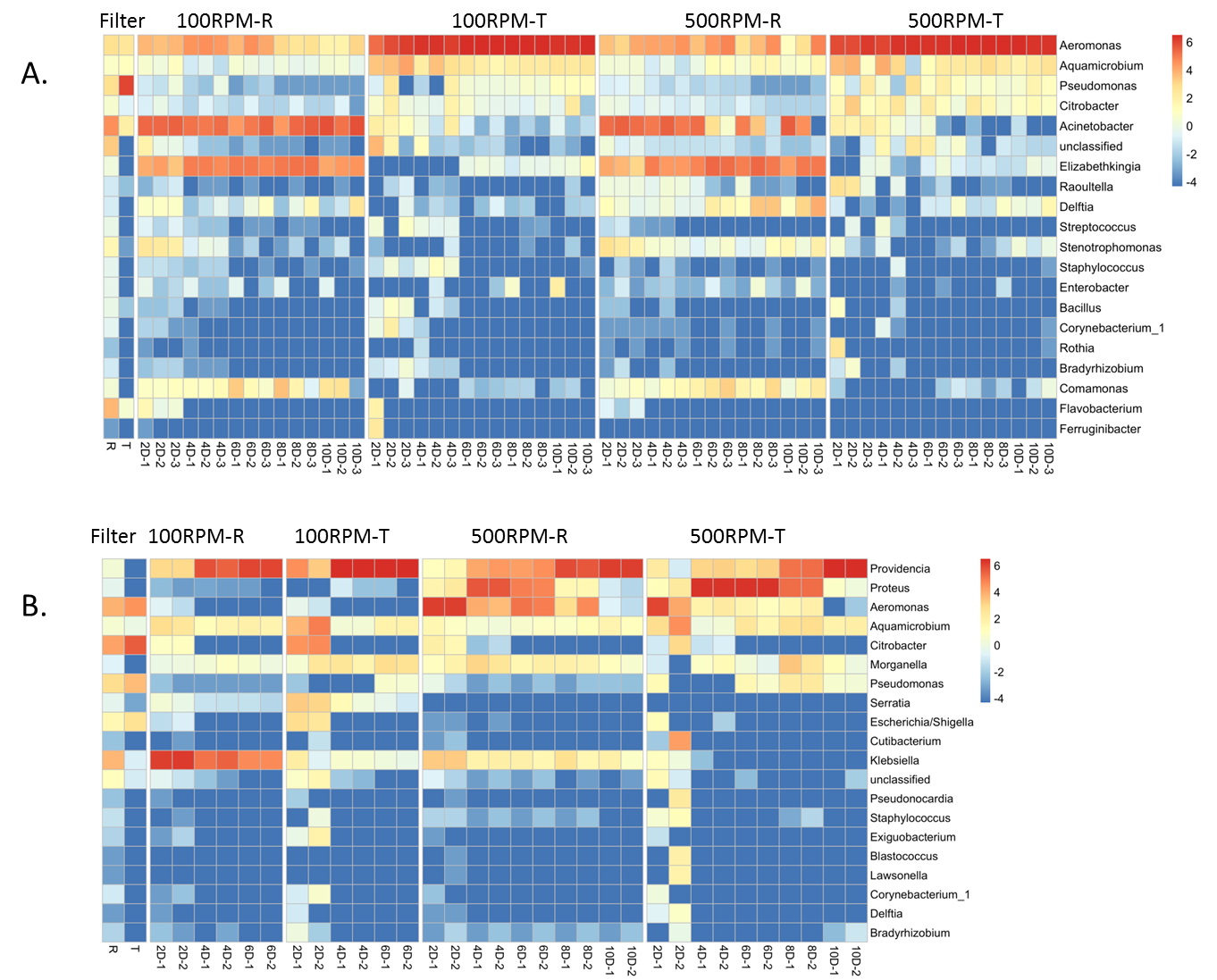

A

B

Figure S7. Relative abundance (log2 transformed) of the 20 most abundant genera in SW (A) and MC (B) media. Filter: filter mating assay, T: transconjugant, R: recipient.

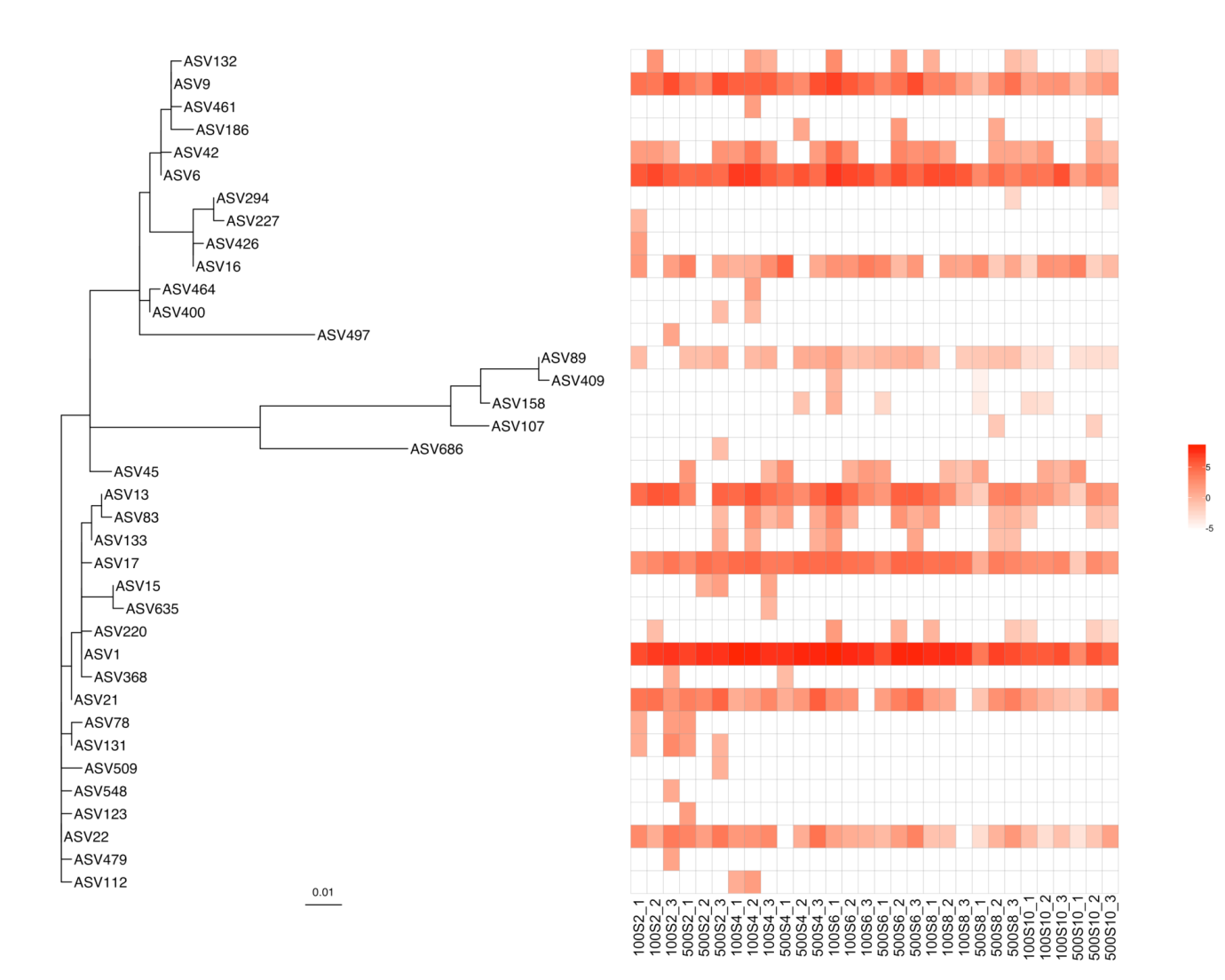

Figure S8. Phylogenetic tree with absolute-abundance heatmap of 37 *Aeromonas* ASVs in serial transfer groups with SW medium.

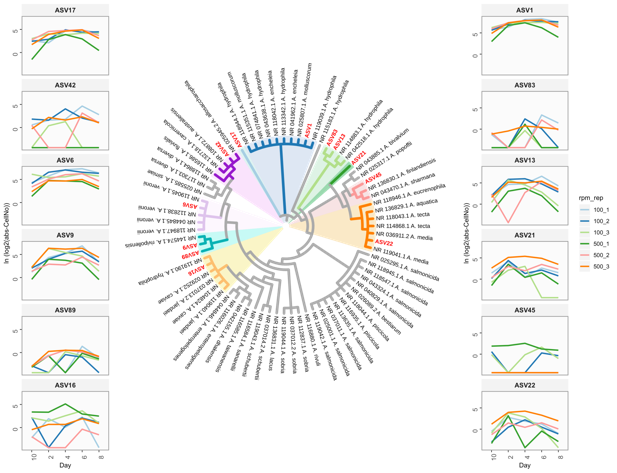

Figure S9. phylogenetic relationship between 12 persistent *Aeromonas* phylotypes (red marked) in SW medium in this study and 63 reference *Aeromonas* sequences from NCBI RefSeq. The sequence clusters with *Aeromonas* phylotypes included were highlighted. The absolute abundance profiles of each of the 12 phylotypes in triplicate serial growth experiments of SW100 and SW500 (indicated by line colour) were shown in plots on two sides of the phylogenetic tree.

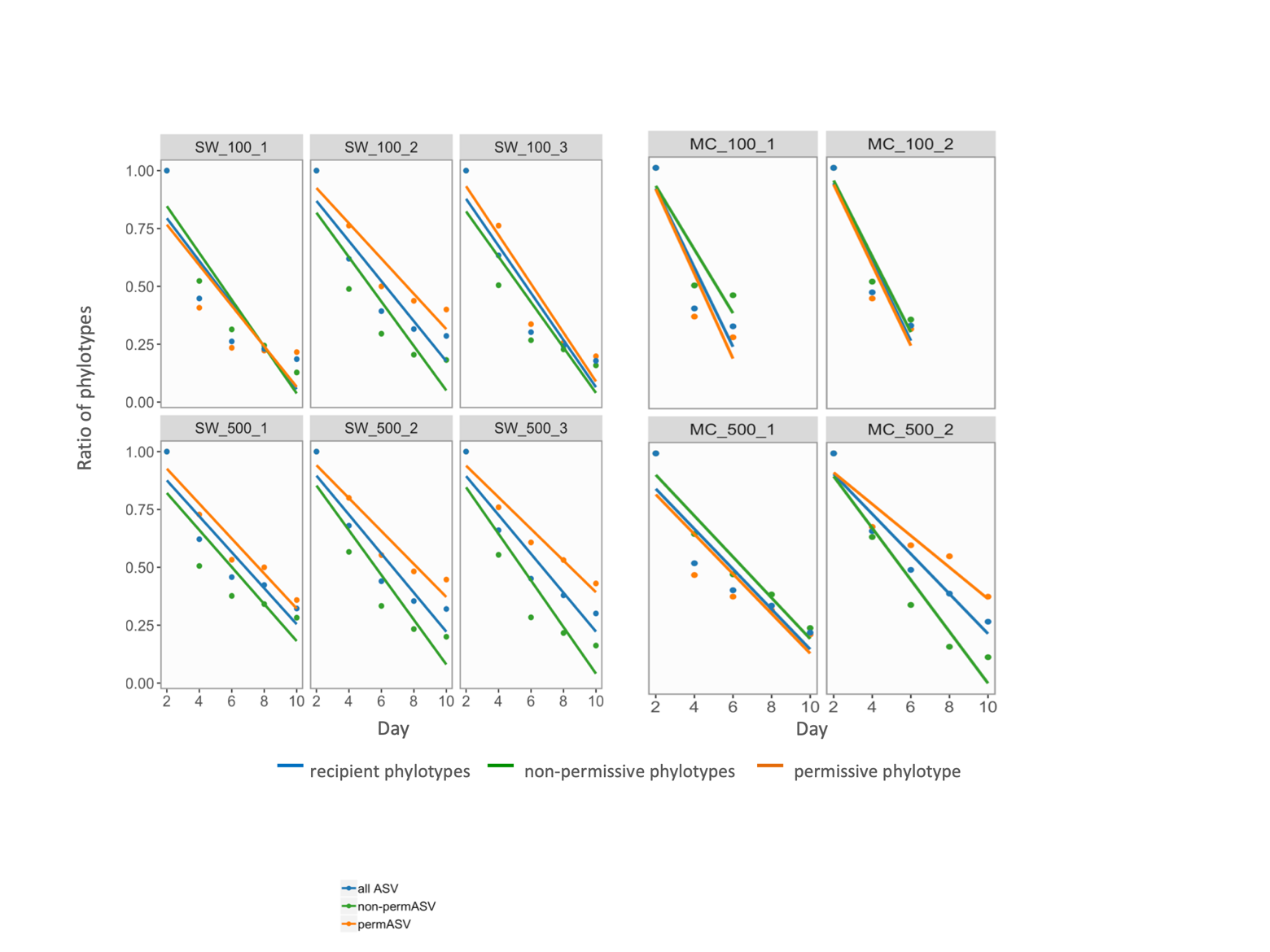

Figure S10. Cumulative loss of phylotypes in recipient pools along serial transfer. X-axis is day of serial transfer. Y-axis is richness of each ASV pool (i.e., recipient phylotypes, non-permissive phylotypes, permissive phylotypes), relative to the first day. Linear regression was used to model loss of unique phylotypes as function of time.

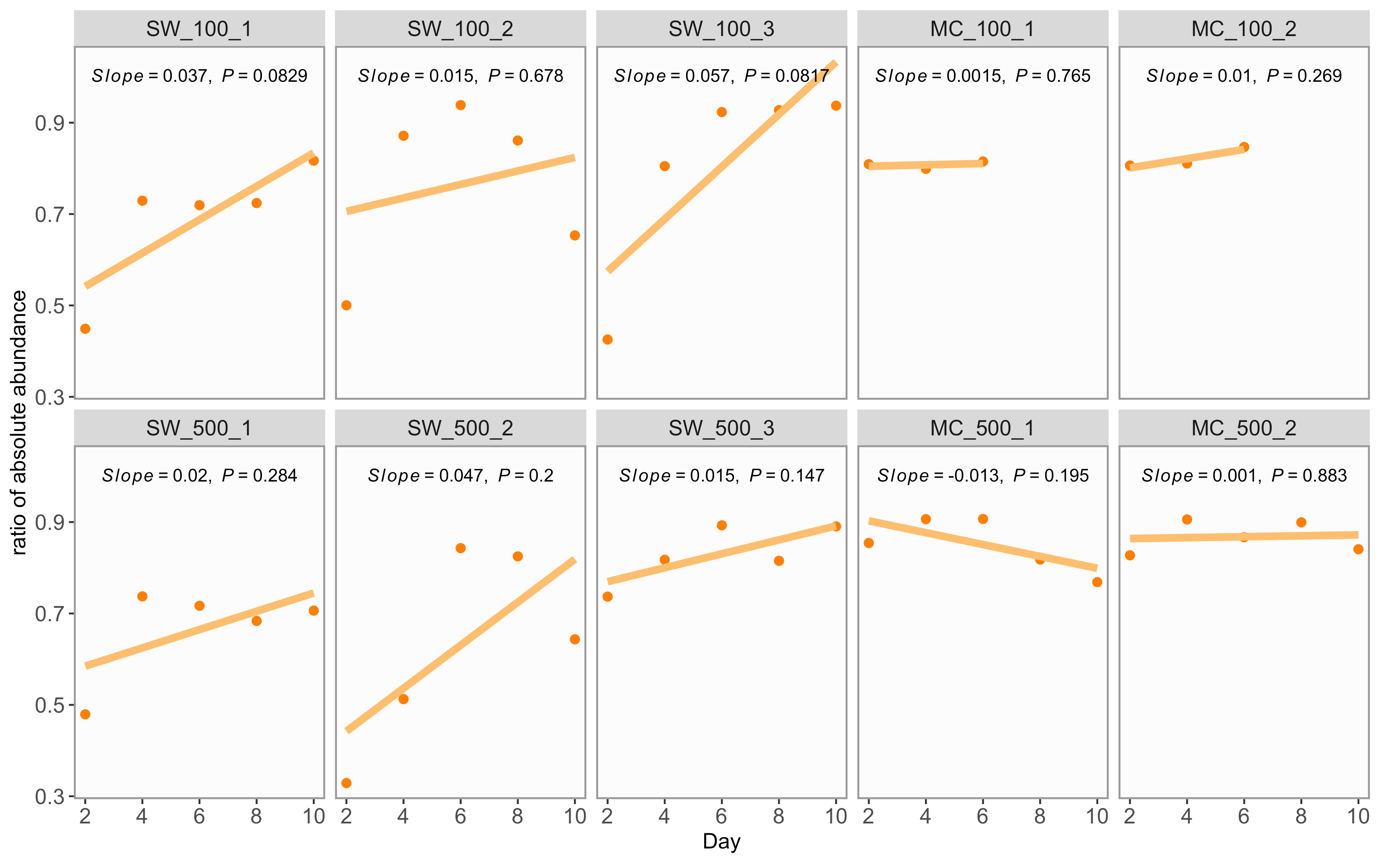

Figure S11. Fitness of permissive phylotypes relative to recipient phylotypes (i.e., time-series change of absolute abundance ratio ($\frac{total abs. abundance of permissve phylotypes in recipient pool}{total abs. abundance of recipient pool}$)). Linear regression model was applied to estimate relative fitness which is trend of absolute abundance ratio as function of time. Solid lines represent regression curve labelled with slope coefficient and P-value. Positive or negative slope indicates overall positive or negative fitness of permissive phylotypes relative to recipient phylotypes. P-value < 0.01 indicated significance of linear correlation between the relative fitness and time series.
